# Hoxb5 Enriches Long-Term Hematopoietic Stem Cell Activity within the Mouse Fetal Liver Phenotypic HSC Compartment

**DOI:** 10.64898/2026.06.30.734854

**Authors:** Victoria L. Mascetti, Allison Banuelos, Kristine Teague, Tuva Wegnelius Jarlstedt, Adam Wilkinson, Hiro Nakauchi, Irving L. Weissman

## Abstract

Hematopoietic stem cells (HSCs) in the adult mouse can be prospectively isolated to near-purity through phenotypic markers, enabling detailed analysis of stem cell function. Homeobox B5 (Hoxb5) was previously identified as a definitive marker of long-term (LT) HSCs in adult bone marrow^1^. In contrast, fetal HSCs have not been purified to the same extent. Here, we show that Hoxb5 is expressed in fetal liver (FL) HSCs at embryonic day (E) 12.5-16.5 using a single-color tri-mCherry reporter driven by endogenous Hoxb5 regulation. Prospective purification by stringent multiparameter flow cytometry revealed Hoxb5⁺ FL-HSCs to exhibit robust, multilineage reconstitution upon serial transplantation. Quantitative assays reveal that Hoxb5 enriches FL-HSCs to near-single-cell purity, analogous to its role in the adult bone marrow, underscoring its reliability in distinguishing LT-HSCs throughout hematopoietic ontogeny. Notably, Hoxb5 expression is not exclusive to FL-HSCs, as it is also detected across the fetal liver hematopoietic hierarchy and in fetal liver endothelial cells, suggesting developmental stage–specific regulation of its expression. In addition, single-cell RNA sequencing of FL-HSCs identified distinct transcriptional states defined by Hoxb5 expression. These findings establish Hoxb5 as a robust marker for enhancing the purification of fetal liver phenotypic HSCs (pHSC) and provide a framework for dissecting the molecular regulation of HSC ontogeny.

## Introduction

Hematopoietic stem cells (HSCs) occupy the apex of the hematopoietic lineage hierarchy and are uniquely defined by their dual capacity for long-term self-renewal and multilineage differentiation. These properties ensure lifelong production of all blood and immune cell types. The ability to prospectively isolate and functionally define HSCs, pioneered in mice in 1988 ^2^ and in humans in 1992 ^3^, has enabled decades of mechanistic insight into stem cell biology and provided the cellular basis for hematopoietic stem cell transplantation in malignant and non-malignant blood disorders. Despite their central role in maintaining hematopoiesis, HSCs are not functionally uniform but exhibit lineage bias, proliferative differences, and age-dependent shifts in output, underscoring the importance of context and developmental stage in defining stem cell potential.

The fetal liver serves as the major hematopoietic organ of mid-gestation, supporting rapid HSC expansion as well as the generation of lymphoid and myeloid progenitors ^4–8^. Fetal liver HSCs are functionally distinct from their adult bone marrow counterparts: they display higher proliferative capacity ^9^, enhanced self-renewal potential ^10^, more long-term repopulating cells relative to short-term repopulating cells ^11^, and broader developmental plasticity ^12^, including the ability to generate specialized Vy3^+^ and Vy4^+^ T cell subsets under the fetal thymic microenvironment ^5^. In humans, though more numerous, aged HSCs are less active than the young HSCs , and as such, the ability of human HSCs to generate primitive progenitors in vitro is higher in the fetal liver compared to umbilical cord blood and is further diminished when HSCs are isolated from adult BM ^13^. These properties make fetal liver HSCs a unique developmental stage for interrogating stem cell fate, plasticity, and long-term regenerative potential.

Much of our knowledge of HSC biology, including evidence for lineage bias and diversity, is deduced from experiments that involve the prospective isolation of HSCs by cell surface marker expression and their transplantation into lethally irradiated mice ^14,15^. It has been shown by ourselves and others that the murine and human hematopoietic cell surface is dramatically reengineered during differentiation, allowing for purification of cell populations by single-cell sorting in conjunction with a panel of fluorescent antibodies ^13^. Efforts to purify fetal liver HSCs have revealed substantial advances yet remain incomplete when compared to adult bone marrow. Early studies demonstrated enrichment of long-term multilineage reconstituting activity within Thy^lo^Lineage^-/lo^Sca-1^+^ fraction ^5,16^, further refined by markers including Mac-1+, CD4- ^7^, and SLAM family receptors (CD150^+^, CD48^-^, CD244^-^) ^17^. Despite these advances, fetal liver phenotypic HSCs (pHSCs) have not yet been prospectively isolated to the near-homogeneity achieved for adult bone marrow pHSCs, where as few as a single cell is sufficient for multilineage engraftment ^18^ and HSC self renewal, usually assayed by serial reconstitution ^15^.

The homeobox transcription factor Hoxb5 was identified as a definitive marker of long-term (LT) HSCs in the adult bone marrow ^1^. While its role in marking adult LT-HSCs is now well established ^19^, whether Hoxb5 similarly defines LT-HSCs during stages of development before adulthood remains uncertain. Given the developmental expression of Hoxb5 during organogenesis and specificity in the adult hematopoietic apex, Hoxb5 represents a strong candidate for enhancing the purification of phenotypic fetal liver HSCs. However, its expression across hematopoietic ontogeny has not been systematically examined. Thus, whether Hoxb5 marks bona fide fetal liver LT-HSCs and can enhance their prospective isolation remains an open question. Here, we address this knowledge gap by investigating Hoxb5 expression in fetal liver HSCs and evaluating its utility for their purification and functional definition.

## Results

### Expression of Hoxb5 on fetal liver hematopoietic stem and progenitor cells

To characterize Hoxb5 expression on fetal liver HSCs (FL-HSCs), we utilized the Hoxb5-tri-mCherry reporter mouse ^1^ and performed multiparameter flow cytometry analysis. Taken from previously published reports demonstrating enrichment of long-term multilineage reconstituting activity within Thy^lo^Lineage^-/lo^Sca-1^+^ fraction ^5,16^, further refined by markers including Mac-1^+^, CD4^-7^, and SLAM family receptors ^17^, we employed a highly stringent gating strategy to interrogate the expression of Hoxb5 on fetal liver HSCs: Lin^-^ cKit^+^ Sca^+^ (LKS) CD48^-^ FLK2^-^ CD150^+^ CD11b^+^ (Fig 1A). We find that Hoxb5 is expressed in fetal liver HSCs at embryonic day (E) 12.5, 14.5 and 16.5 (Fig 1B-E). There is an expansion in the absolute number of Hoxb5 HSCs per fetal liver at E14.5 with expression of Hoxb5 in the HSC compartment from E12.5 to E16.5 following a normal rather than linear distribution. To control for developmental stage variance in the size or total number of cells in a fetal liver we show that the Hoxb5^+^ HSC count per 10000 KLS cells also follows the same normal distribution (Fig 1C), confirming an expansion in the number of Hoxb5^+^ HSCs at E14.5. In line with the expansion of the HSC compartment at this stage (E14.5) previously reported ^8,12^, the expanding population of Hoxb5^+^ HSCs at E14.5 would suggest that Hoxb5 represents a new marker of fetal liver HSCs. The distribution and frequency of Hoxb5 expression is also dependent on the purity of the HSPC subset: For E12.5-16.5 fetal liver, the majority of LKS are Hoxb5^neg^ or Hoxb5^lo^ with Hoxb5^+^ representing 13.7-6.4% frequency of the population, this is contrasted to 69.9-39.1% Hoxb5^+^ cells in the HSC compartment (Fig 1F). There is also evident a greater frequency of Hoxb5^hi^ cells in E12.5 HSC compared to later fetal liver stages, which may be indicative of the establishment of Hoxb5^+^ HSCs in the fetal liver niche at this time, likely seeded as HoxB5^+^ pre-HSC from the embryonic stage of development. Similarly, Hoxb5^+^ frequency increases with population purity to be enriched in the HSC compartment [LKS CD48^-^ FLK2^-^ CD150^+^ CD11b^+^] (Fig 1G), with a mean total count per fetal liver of 178 cells at E14.5 (Fig 1H).

**Figure 1.**
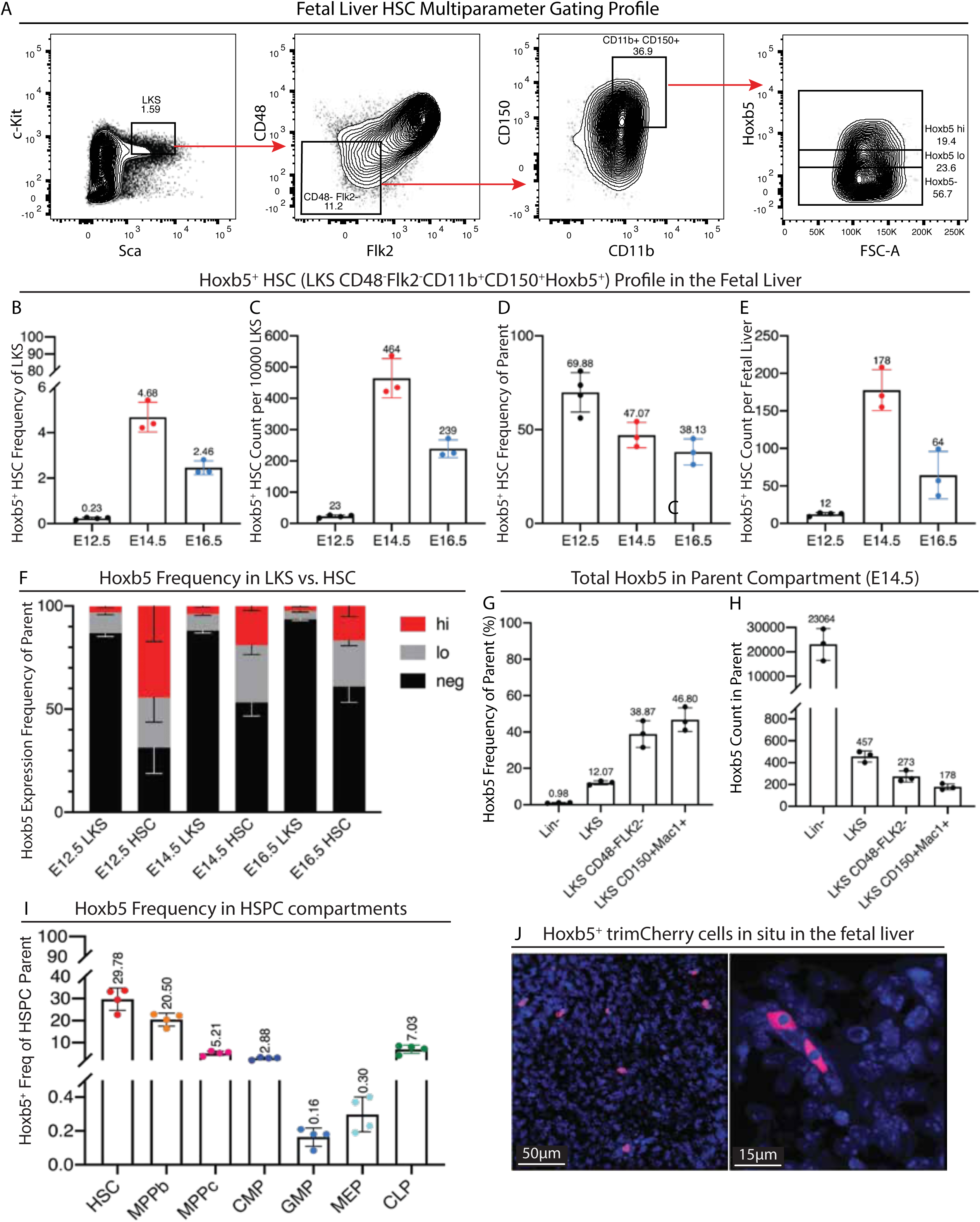
Hoxb5 is expressed on fetal liver hematopoietic stem and progenitor cells. A. Flow cytometry plots of E14.5 fetal liver from mice depicting Hoxb5-tri-mCherry reporter activity in HSC immunophenotype Lin^-^ cKit^+^ Sca^+^ (LKS) CD48^-^ FLK2^-^ CD150^+^ Mac1^+^; B-H. Hoxb5^+^ HSC profile at E12.5, E14.5 and E16.5. B. Hoxb5^+^ HSC frequency of LKS population. C. Hoxb5^+^ HSC count per 10000 LKS. D. Hoxb5^+^ frequency in HSC population (Lin^-^ cKit^+^ Sca^+^ CD48^-^ FLK2^-^ CD150^+^ Mac1^+^). E. Total Hoxb5^+^ HSC count per whole fetal liver. F. Cell frequency of Hoxb5 negative (neg), low (lo), and high (hi) expressing cells in LKS or HSC compartment at E12.5, E14.5 and E16.5. G-H. Hoxb5^+^ cell frequency in each sequential multiparameter gate from Lin- to HSC compartment [Lin^-^, LKS, CD48^-^ FLK2^-^, CD150^+^ Mac1^+^]. I. Hoxb5 frequency in hematopoietic stem and progenitor cell (HSPC) compartments including HSC, MPPb, MPPc, CMP, GMP, MEP, CLP. J. Immunocytochemistry of Hoxb5-tri-mCherry positive cells in situ in the fetal liver.

To test if Hoxb5 provides as exclusive marker of HSCs in the fetal liver we analyzed the expression of Hoxb5 in all sequential multiparameter flow cytometry gates from live cells to HSCs (Supplemental Fig. 1). We found that Hoxb5 is represented in live and lineage negative cells of the fetal liver. However, Hoxb5^+^ cells are represented at progressively greater frequency as sequential gating analysis proceeds and culminates in the HSC subset [Lin^-^ 0.98%; LKS 12.07%; LKS CD48^-^ Flk2^-^38.87%; HSC (LKS CD48^-^ Flk2^-^ CD150^+^ Mac1^+^) 46.80%].

To determine the expression of Hoxb5 in hematopoietic stem and progenitor cell (HSPC) subsets, we performed flow cytometry of fetal livers from mice depicting Hoxb5-tri-mCherry reporter activity in previously defined HSPC immunophenotypes: HSC [Lin^-^ Sca^+^ cKit^+^ CD150^+^ FLK2^-^ CD34^-^]; multipotent progenitor (MPP) b [Lin^-^ Sca^+^ cKit^+^ CD150^-^ FLK2^-^ CD34^+^]; MPPc [Lin^-^ Sca^+^ cKit^+^ CD150^-^ FLK2^+^ CD34^+^]; common myeloid progenitor (CMP) [Lin^-^ Sca^lo/-^ cKit^+^ CD34^med/hi^ CD16/32^-/lo^]; megakaryocytic-erythroid progenitor (MEP) [Lin^-^ Sca^lo/-^ cKit^+^ CD34^-^ CD16/32^-/lo^ CD150^+^]; granulocyte-monocyte progenitor (GMP) [Lin^-^ Sca^lo/-^ cKit^+^ CD34^hi^ CD16/32^hi^] (Fig 1I and Supplemental Figure 5). We find Hoxb5 expressing cells in all hematopoietic stem and progenitor compartments with the greatest frequency of Hoxb5^+^ cells in HSC (29.78%) and MPPb (20.50%) compartments. Therefore, Hoxb5 does not serve as an exclusive marker of fetal liver HSCs but rather an additional marker to enhance the purity of HSC isolation.

### Hoxb5 enhances phenotypic HSC reconstitution activity

Prospective isolation of pHSCs requires that the isolated cells are capable of long-term production of all blood cells in primary irradiated hosts, as well as self-renewal, such that the cells can be transplanted to secondary hosts to give rise to long-term multilineage reconstitution ^15^. To determine the phenotypic nature of the Hoxb5 fraction of fetal liver HSCs [LKS CD48^-^ FLK2^-^ CD150^+^ CD11b^+^ Hoxb5^+/-^], we performed competitive transplant of sorted Hoxb5^+^ or Hoxb5^-^ E14.5 fetal liver HSCs (CD45.2^+^) with CD45.1^+^ support cells into lethally irradiated adult recipients (CD45.1^+^) (Fig. 2A). Analysis of Hoxb5^+^ donor peripheral-blood chimerism 16 weeks after transplant demonstrated multilineage reconstitution in both myeloid (GR1^+^ granulocytes and Mac1^+^ monocytes) and lymphoid (CD3^+^ T cells and B220^+^ B cells) lineages by donor CD45.2^+^ fetal liver HSCs (Fig 2B). At limiting dilution of 200, 100, 10, and 1 cell, Hoxb5^+^ fetal liver HSCs contributed robustly to both myeloid and lymphoid lineages (Fig 2C, 2E), with durable engraftment observed from 4 to 16 weeks post-transplant (Fig. S2). Importantly, secondary transplantation of whole bone marrow from primary recipients demonstrated that Hoxb5^+^ fetal liver HSCs retained long-term, serial repopulating capacity, establishing them as bona fide LT-HSCs (Fig. 2D, 2F).

**Figure 2.**
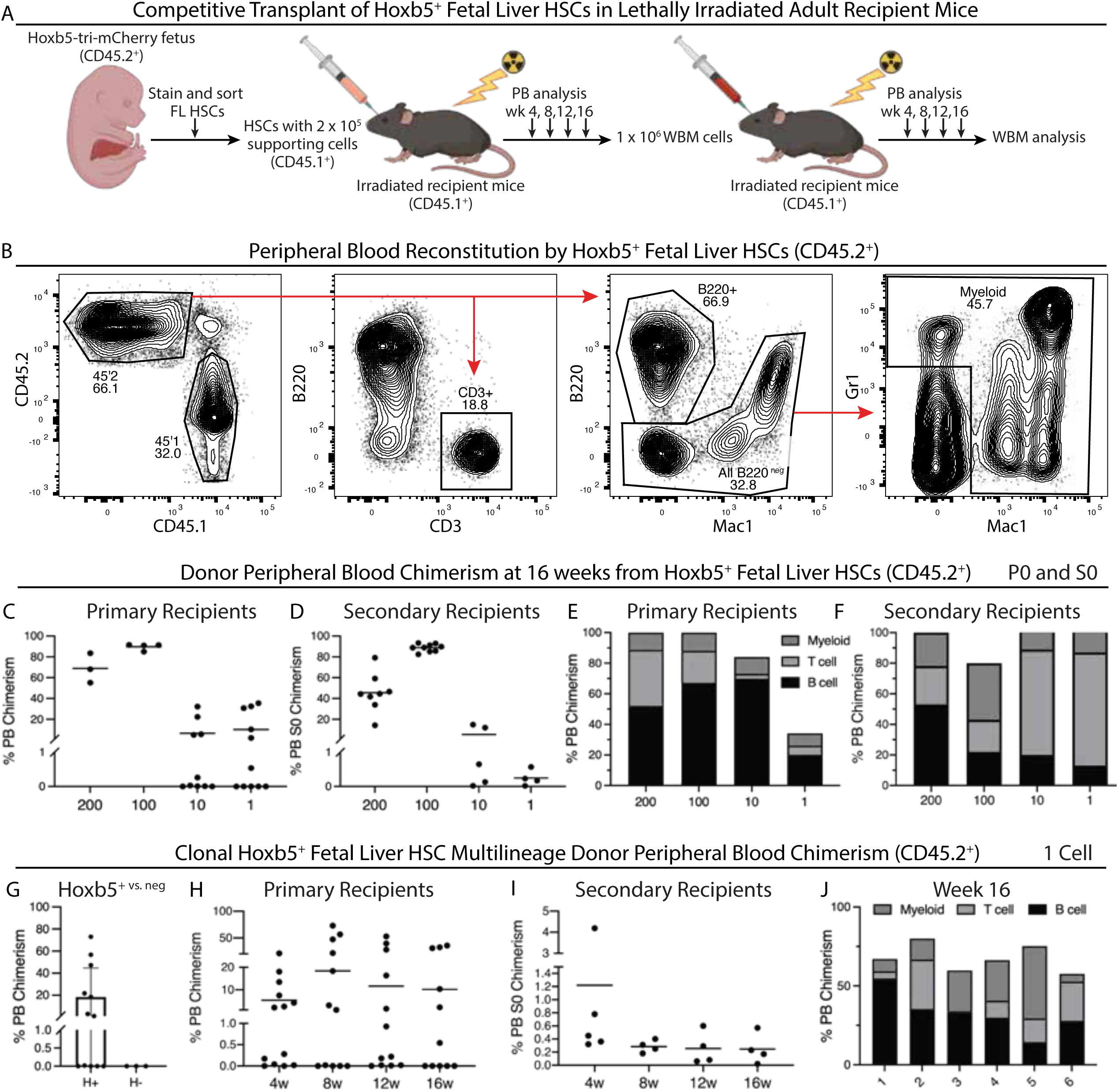
Hoxb5 distinguishes the long-term reconstituting HSC fraction in the fetal liver. A. Experimental schematic for long-term haematopoietic reconstitution by competitive transplant. Hoxb5–tri-mCherry CD45.2^+^ HSCs (200, 100, 10, or 1 cells) and 2x10^5^ CD45.1^+^ supporting cells were competitively transplanted to lethally irradiated CD45.1^+^ recipient mice. For secondary transplants, 1x10^6^ whole bone marrow (WBM) cells were transferred from primary recipient mice into lethally irradiated secondary recipient mice. PB, peripheral blood. B. Donor peripheral-blood chimerism FACS gating scheme for CD45.2^+^-derived lymphoid (CD3^+^ T cells and B220^+^ B cells) and myeloid lineages (GR1^+^ granulocytes and Mac1^+^ monocytes). C-F. Multilineage donor peripheral-blood chimerism at 16 weeks for each individual mouse from 200, 100, 10, or 1 Hoxb5^+^ FL-HSCs (CD45.2^+^). Peripheral blood chimerism in primary recipients (C), secondary recipients (D), mean peripheral blood donor chimerism by HSC-derived myeloid or lymphoid (T cell and B cell) lineages in primary (E) and secondary (F) recipients. G. Clonal multilineage donor peripheral blood chimerism (CD45.2^+^) after single cell competitive reconstitution of Hobx5^+^ vs Hoxb5^-^ single (1 cell) FL-HSC. H-J. Clonal donor peripheral blood chimerism at week 4-16 in primary (H) and secondary (I) recipients. J. Week 16 multilineage peripheral blood reconstitution for each individual secondary recipient mouse following clonal reconstitution of the primary recipient.

Mice transplanted with as many as 10 Hoxb5⁻ fetal liver HSCs [[LKS CD48^-^ FLK2^-^ CD150^+^ CD11b^+^ Hoxb5⁻] failed to achieve durable hematopoietic reconstitution and succumbed between 8–12 weeks post-transplant. Although all recipients were co-transplanted with CD45.1⁺ whole bone marrow support cells to provide short-term radioprotection, these cells lack long-term self-renewing capacity and therefore cannot sustain hematopoiesis beyond the early post-transplant period. The observed mortality in the absence of engraftment by the Hoxb5⁻ phenotypic HSC fraction demonstrates that long-term repopulating activity is restricted to the Hoxb5⁺ fraction within the phenotypically defined HSC compartment. This is consistent with prior studies demonstrating that committed progenitors provide only transient survival following lethal irradiation ^15,20,21^.

Peripheral blood analysis revealed as few as a single Hoxb5^+^ fetal liver HSC was sufficient to generate multilineage chimerism, whereas Hoxb5^-^ fetal liver HSC lacked durable contribution (Fig. 2G-2J). Single-cell transplantation of a Hoxb5⁺ fetal liver HSC demonstrated that 6 of 11 recipients achieved multilineage reconstitution, corresponding to an engraftment success rate 55% and probability of 0.55 (95% CI: 0.23–1.83). Fitting these data to a single-hit Poisson model yielded an estimated stem cell frequency of 1 in 1.27 cells (95% CI: 1 in 0.6 to 1 in 3.8). When compared to previously published prospective isolation using SLAM family markers [LKS CD48^-^ FLK2^-^ CD150^+^ CD48 CD11b^+^] 1 in 2.7 cells (37%) gave competitive reconstitution of lethally irradiated mice, this reveals that Hoxb5 enhances the purity of fetal liver LT-HSCs. These findings parallel the function of Hoxb5 in adult bone marrow, where it prospectively identifies LT-HSCs ^1^, and extend its utility to developmental hematopoiesis by demonstrating that Hoxb5 enriches fetal liver LT-HSCs to near-single-cell purity.

### Hoxb5^+^ fetal liver HSCs reconstitute adult bone marrow in recipient mice

To assess whether Hoxb5^+^ fetal liver HSCs give rise to bona fide LT-HSCs within the adult bone marrow niche, we analyzed bone marrow HSC [lin^-^ CD34^-^ cKit^+^ Sca1^+^] reconstitution in primary and secondary lethally irradiated recipients (CD45.1^+^) transplanted with 200, 100, 10, or 1 Hoxb5^+^ fetal liver HSCs (CD45.2^+^) and competitor support cells (CD45.1^+^) (Fig. 3A). At 18 weeks post-transplant, donor-derived CD45.2^+^ cells dominated total bone marrow chimerism following 200 and 100 cell transplant, whereas host-derived CD45.1^+^ cells contributed minimally, indicating durable engraftment of transplanted Hoxb5^+^ fetal liver HSCs (Fig. 3B). For 10 and single Hoxb5+ fetal liver HSC competitive transplants, chimerism in the adult bone marrow was also considered well established (10 cell: 11.37-17.99% CD45.2^+^, 1 cell: 4.63-19.89% CD45.2^+^). Frequency of clonal bone marrow reconstitution by a single CD45.2^+^ donor derived fetal liver LT-HSC is shown (Fig 3C). Within the hematopoietic stem and progenitor compartment, CD45.2^+^ cells were highly enriched in the Lineage^-^c-Kit^+^ Sca-1^+^ (LKS) fraction, consistent with robust self-renewal and maintenance of the stem cell pool (Fig 3C). Strikingly, analysis of Hoxb5 expression revealed that reconstituted LT-HSCs within the bone marrow were almost exclusively of donor (CD45.2^+^) origin, with negligible contribution from competitor (CD45.1^+^) cells. These results establish that Hoxb5^+^ fetal liver LT-HSCs not only sustain long-term multilineage hematopoiesis, but also generate a self-renewing population of HSCs in the adult bone marrow, mirroring the defining properties of adult LT-HSCs ^1,15,22^.

**Figure 3.**
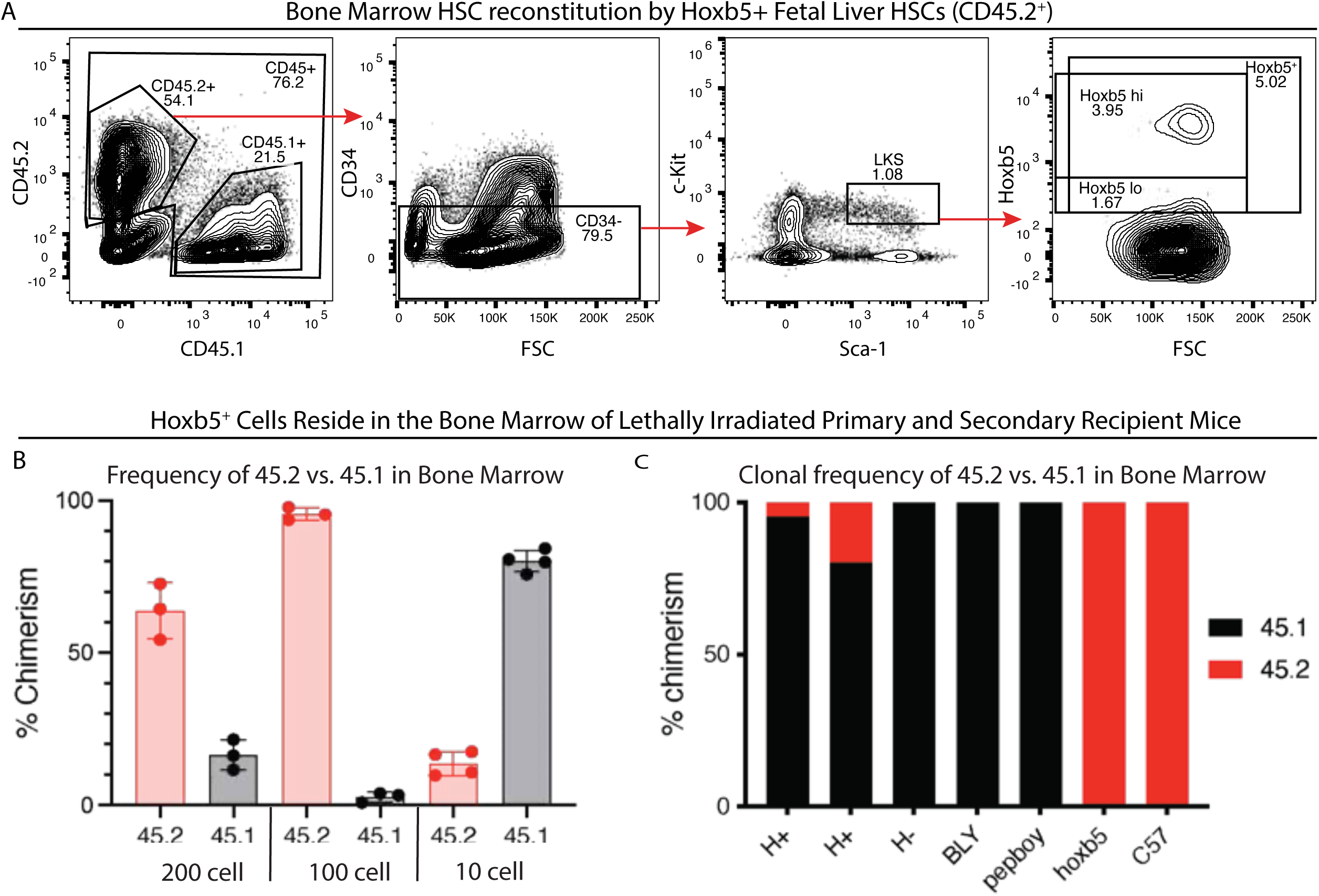
Hoxb5+ FL-HSC reconstitute the bone marrow of lethally irradiated recipients. A. Flow cytometry plots of bone marrow from 16-week CD45.1^+^ recipient mice depicting donor-derived the CD45.2^+^ fraction of immunophenotypic HSCs (Lin^-^, CD34^-^ c-kit^+^, Sca1^+^) with Hoxb5 expression shown. B. Frequency of CD45.2^+^ or CD45.1^+^ in the bone marrow of reconstituted CD45.1^+^ recipients following transplants with 200, 100, or 10 CD45.2^+^ Hoxb5^+^ FL-HSCs. C. Bone marrow chimerism by a single donor CD45.2^+^ Hoxb5^+^ FL-HSC compared with Hoxb5^-^ Fl-HSC, included are non-irradiated controls for CD45.1 BLY and CD45.1 Pepboy recipient backgrounds, and donor CD45.2 Hoxb5 Hoxb5–tri-mCherry reporter mouse and CD45.2 C57 mouse.

### Hoxb5 expression in fetal liver HSCs defines distinct transcriptional states

To define the transcriptional features underlying Hoxb5 expression in fetal liver HSCs and progenitor cells (HSPCs), we performed single-cell RNA sequencing (scRNA-seq) of FACS-purified fetal liver cells from E12.5, E14.5, and E16.5 Hoxb5-tri-mCherry reporter mice. Unsupervised clustering of the combined dataset revealed transcriptionally distinct populations corresponding to hematopoietic stem cells (HSCs), multipotent progenitors (HSPCs), and lineage-committed progenitors (CMP, GMP, MEP) (Fig. 4A). Within the HSC cluster Hoxb5 m-Cherry reporter status (Fig. 4B) and Hoxb5 RNA expression was visualized (Fig. 4C). Two major transcriptional HSC clusters were identified: Cluster 1 enriched for Hoxb5⁺ fetal liver HSCs, and cluster 2 enriched for Hoxb5⁻ fetal liver HSCs, representing transcriptionally distinct stem and progenitor states (Fig. 4D–H, Supplemental Figure 3). Differential gene expression analysis between HSC clusters 1 and 2 demonstrated that Cluster 1 enriched for Hoxb5⁺ HSCs express high levels of canonical stemness genes including Hlf, Mecom, Fgd5, and Mllt3, consistent with a quiescent, self-renewing phenotype (Fig. 4I). The enrichment of Mllt3 in Hoxb5+ fetal liver HSCs is notable given that MLLT3 has been identified as a conserved regulator of human HSC self-renewal and engraftment ^23^. In contrast, Cluster 2 enriched for Hoxb5⁻ HSCs exhibited elevated expression of proliferation- and differentiation-associated genes such as Mpo, Tespa1, and CD48 suggesting a transition toward progenitor identity (Fig. 4I).

**Figure 4.**
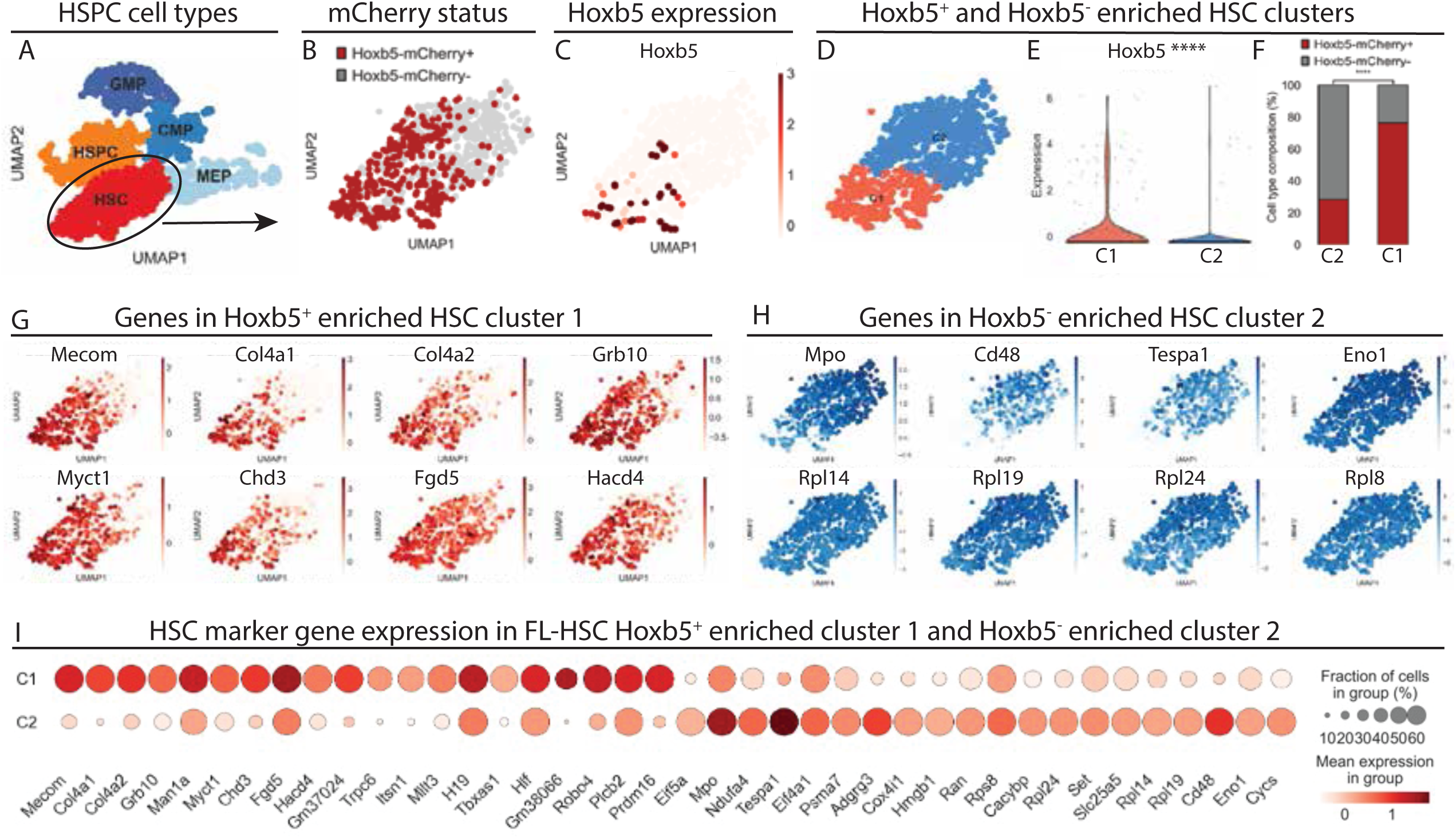
Hoxb5 marks a transcriptionally distinct subset of fetal liver HSCs. **A.** Single-cell RNA sequencing of index-sorted HSPCs from E12.5, E14.5, and E16.5 fetal livers of Hoxb5-mCherry reporter mice using Smart-seq3. Plot is a UMAP-embedded representation of Leiden clusters annotated by cell type identity based on expression of known lineage marker genes. B. UMAP highlighting Hoxb5-mCherry reporter status within the HSC cluster, as determined by index-sorting information (red, Hoxb5-mCherry⁺; gray, Hoxb5-mCherry⁻). C. RNA expression of Hoxb5 across the HSC cluster visualized by expression intensity. D. Unsupervised subclustering of HSCs reveals two transcriptionally distinct subsets, cluster 1 (C1) and cluster 2 (C2). E. Violin plot showing Hoxb5 RNA expression in C1 and C2 (Mann–Whitney U = 89 198.00, p = 2.46 × 10⁻¹⁵). F. Fraction of Hoxb5-mCherry⁺ cells in C1 and C2. Association between Hoxb5 status and cluster identity was assessed by chi-square test (χ² = 173.21; p = 1.47 × 10⁻³⁹) and Fisher’s exact test (odds ratio = 8.21; p = 7.52 × 10⁻⁴¹). G-H. Feature plots showing representative genes enriched in C1 (Hoxb5⁺ HSCs; G) and C2 (Hoxb5⁻ HSCs; H). I. Dot plot showing the top 20 genes upregulated in Hoxb5⁺ (C1) and Hoxb5⁻ (C2) fetal liver HSCs.

Analysis of fetal liver hematopoietic stem and progenitor cell (HSPC) subsets confirmed that Hoxb5 expression marks transcriptionally distinct stem and progenitor fractions (Supplemental Fig. 6). Within HSPC subsets, Hoxb5, Hlf, and Mecom were co-expressed at highest levels in the HSC population, at lower levels in MPPs, and were largely absent in committed progenitors (CMP, GMP, and MEP) (Supplemental Fig. 6). This gradient of expression mirrors the phenotypic enrichment of Hoxb5 observed by flow cytometry and supports a role for Hoxb5 in defining the transcriptional identity of long-term fetal-liver HSCs.

### Temporal modulation of fetal liver HSC transcriptional dynamics and Hoxb5 expression

Integration of datasets across these developmental stages and Hoxb5 status revealed that fetal liver HSCs form a continuous but temporally structured transcriptional trajectory, reflecting coordinated shifts in gene expression as the fetal liver transitions from a site of HSC expansion to one of progenitor differentiation (Fig 5A-C). Unsupervised clustering and pseudotemporal alignment identified stage-specific transcriptional profiles associated with early (E12.5), mid- (E14.5), and late (E16.5) fetal liver development (Fig 5D-G). E12.5 fetal liver HSCs exhibited enriched expression of genes such as Runx1, and CD34 consistent with but not proof of their proposed recent emergence from hemogenic endothelium ^24,25^. These cells also displayed high expression of Hlf, Mecom, and Procr, canonical markers of primitive, quiescent HSCs. By E14.5, the FL-HSC transcriptional landscape shifted toward genes associated with self-renewal and metabolic activity, including Mllt3 and Kit, reflecting the well-characterized proliferative expansion of the fetal HSC pool at this stage ^7,8,12,23^. This period corresponds to peak HSC expansion in the fetal liver niche, proposed to be supported by hepatoblast-, endothelial-, and macrophage-derived growth factors ^26,27^. At E16.5, FL-HSCs began to express transcriptional signatures associated with lineage priming, including upregulation of Cd48, Mpl, and Flt3, together with gradual downregulation of Mecom and Slamf1. This shift reflects the onset of differentiation and migration toward the bone marrow niche ^7,11^.

**Figure 5.**
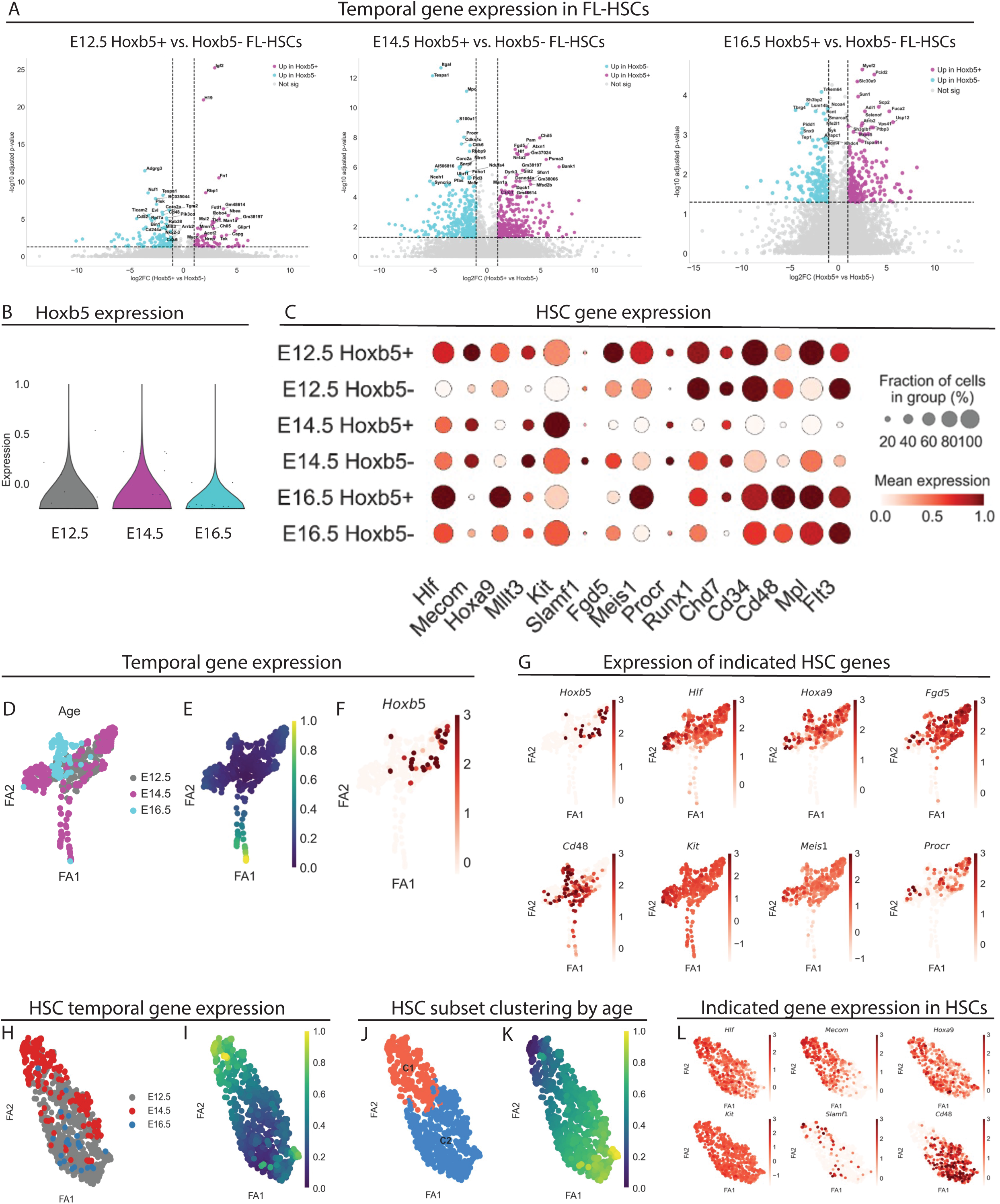
Temporal dynamics of Hoxb5⁺ and Hoxb5⁻ fetal liver HSCs. A. Differential gene expression analysis of Hoxb5⁺ and Hoxb5⁻ fetal liver HSCs at E12.5, E14.5, and E16.5. Volcano plots show upregulated and downregulated genes in Hoxb5⁺ versus Hoxb5⁻ HSCs at each developmental stage. B. Violin plot showing Hoxb5 RNA expression levels in fetal liver HSCs across E12.5, E14.5, and E16.5. C. Dot plot summarizing expression of canonical HSC genes (Hlf, Mecom, Hoxa9, Mllt3, Kit, Slamf1, Fgd5, Meis1, Procr, Runx1, Cd34, Flt3, and others) in Hoxb5⁺ and Hoxb5⁻ HSCs. D-F. Force-directed layout (FA) of all fetal liver HSCs colored by developmental stage (D) and diffusion pseudotime (E), Hoxb5 expression (F), showing a continuous transcriptional trajectory from E12.5 to E16.5. G. Feature plots showing expression of key HSC genes. H-I. FA embedding of HSCs colored by developmental stage (H) and pseudotime (I), showing coordinated transcriptional maturation from E12.5 to E16.5. J-K. FA embedding of HSCs colored by C1 and C2 leiden clustering (J) and corresponding pseudotime (K). L. Feature plots showing expression of key HSC genes.

Longitudinal analysis of Hoxb5 expression in fetal liver HSCs revealed that it is dynamically regulated across development, peaking at E14.5 (Fig. 5B), aligning with both the numerical and functional expansion of reconstituting HSCs shown by flow cytometry and transplantation (Fig. 1–3). This pattern parallels the transition from a proliferative fetal niche to a quiescent adult-like bone marrow environment and is consistent with findings that Hoxb5 marks self-renewing LT-HSCs in adult settings ^1,19^. Unsupervised clustering of fetal liver HSCs across developmental stages identified two transcriptionally distinct HSC subsets, designated cluster 1 [Hoxb5^+^-enriched] and cluster 2 [Hoxb5^-^-enriched] (Fig. 4D-F, Fig. 5J). While pseudotime trajectory analysis placed cluster 1 HSCs at the root of the developmental continuum, giving rise to cluster 2 as an intermediate state preceding differentiation (Fig. 5K), this methodology can reveal markers for physiological lineage tracing, but it is still a morphology analysis. This proposed progression from cluster 1 to cluster 2 could capture the early transcriptional transition from a long-term Hoxb5⁺ fetal liver HSC toward a more metabolically active, Hoxb5⁻ progenitor state within the fetal liver niche.

## Discussion

Our findings establish Hoxb5 as a robust marker of fetal liver LT-HSCs, extending its previously defined role in adult bone marrow ^1^ to the earliest serially transplantable stage of hematopoietic ontogeny. By using an endogenous Hoxb5 reporter combined with stringent multiparameter flow cytometry, we demonstrate that Hoxb5^+^ fetal liver HSCs are capable of multilineage, long-term reconstitution at the single-cell level and retain serial transplantation capacity as definitive hallmarks of bona fide LT-HSCs ^15,22^. These data place Hoxb5 alongside the most powerful markers for HSC purification to date and provide a new tool for probing stem cell function during fetal development.

The ability to prospectively enrich fetal liver HSCs has been the focus of multiple decades of study. Early enrichment strategies used Thy^lo^Lineage^-/lo^Sca-1^+^ fractions ^5,16^, later refined by Mac-1, CD4 ^7^, and SLAM family receptors (CD150^+^, CD48^-^, CD244^-^) ^17^. More recent approaches, including transcriptional profiling, have underscored the heterogeneity of fetal liver HSCs and revealed subsets with distinct self-renewal and lineage output. Comparison of reconstitution efficiency between Hoxb5^+^ fetal liver pHSCs and prior strategies indicate that Hoxb5 enhances the purity of fetal liver LT-HSCs to near-single-cell resolution, analogous to its role in the adult ^1^.

Functionally, fetal liver HSCs are distinct from adult bone marrow HSCs in their higher proliferative capacity, enhanced self-renewal, and broader lineage outcomes that could be a sign of plasticity ^9,10,12^ or distinct HSC subsets that don’t interconvert ^28^. Recent clonal analyses have further demonstrated that hematopoietic output is intrinsically heterogeneous even under native conditions ^29^. Our finding that Hoxb5 enriches for the long-term repopulating fraction across ontogeny indicates that it identifies a conserved stem cell state shared between developmental and adult stages.

RNA-seq studies of fetal liver hematopoietic compartments ^26,27,30,31^ reveal transcriptional programs distinct from adult HSCs populations, including proliferative and metabolic signatures. Elsewhere we find that the neogenin 1+ fetal liver HSC are myeloid biased in outcomes and self-renewal, while the neogenin 1- subsets are balanced in outcomes and self-renewal (Banuelos et al, in preparation). Mapping Hoxb5 expression within these contexts will aid in dissecting the molecular regulation of HSC emergence, including defining independent HSC subsets or lineage relationships, and may facilitate efforts to generate functional HSCs in vitro from pluripotent stem cells. Collectively, these single-cell data demonstrate that Hoxb5⁺ fetal-liver HSCs represent a transcriptionally distinct and developmentally stable stem cell state enriched for self-renewal and quiescence programs, while Hoxb5⁻ cells show progressive differentiation and reduced stemness, similar to short-term adult bone marrow HSCs ^15^. This molecular segregation aligns with the functional data demonstrating that only the Hoxb5⁺ fraction sustains long-term, multilineage hematopoietic reconstitution following transplantation. The temporal dynamics observed here are consistent with recent human and mouse single-cell studies demonstrating stage-dependent changes in fetal hematopoietic programs ^26,27,30,31^. Our data extend these findings by providing a high-resolution view of likely Hoxb5-regulated transcriptional trajectories within fetal liver HSCs.

Notably, Hoxb5 expression is not exclusive to HSCs: we observe expression across progenitor subsets and within fetal liver endothelial cells. This is consistent with the developmental history of Hoxb5, originally identified as Hox2.1 in embryonic organogenesis ^32^, where it plays roles in mesodermal and endothelial patterning. Our observation that Hoxb5 is detectable in endothelial subsets (data not shown) but markedly enriched in phenotypically defined HSCs suggests that its regulation may reflect the proposed developmental continuum linking vascular endothelium and definitive hematopoiesis ^24,33,34^. This interpretation is consistent with studies demonstrating that definitive HSCs emerge within specialized embryonic vascular niches, including the placental vasculature ^35,36^, before subsequently undergoing extensive expansion within the fetal liver ^7,12^. It may also reflect the shared transcriptional program observed between adult bone marrow HSCs and endothelial cells, including expression of genes such as Runx1, Tie2 (Tek), VE cadherin (Cdh5), CD31 (Pecam1) ^22,34,37–41^. This dual representation highlights the importance of contextual marker panels: while Hoxb5 is not sufficient alone, in combination with established HSC markers it enables highly stringent prospective isolation. Although Hoxb5 expression was detected across multiple fetal liver HSPC compartments, competitive primary and serial transplantation demonstrated that durable, multilineage, self-renewing activity was restricted to the Hoxb5⁺ fraction within the phenotypically defined HSC compartment. We did not evaluate the functional potential of Hoxb5⁺ cells within downstream progenitor populations and therefore do not infer equivalent stem cell activity based solely on shared Hoxb5 expression. Rather, our findings indicate that Hoxb5 functions as a marker that enriches for long-term repopulating activity specifically within the phenotypically defined fetal liver HSC population.

In conclusion, this study demonstrates that Hoxb5 expression enriches fetal liver LT-HSCs with high fidelity, aligning with its established role in the adult. While not exclusive to HSCs, Hoxb5 provides a powerful handle for prospective isolation, testing the proposed endothelial–hematopoietic transition and enabling dissection of stem cell regulation across ontogeny. These insights refine our understanding of fetal liver HSC biology and provide a framework for harnessing developmental programs to engineer transplantable HSCs.

## Resource availability

Requests for further information and resources should be directed to and will be fulfilled by the lead contact, Dr Victoria L. Mascetti. This study did not generate new unique reagents. Single-cell RNA-seq data have been deposited at GEO and are publicly available as of the date of publication.

## Acknowledgments

We would like to acknowledge T. Naik and T. Storm for laboratory management; A.McCarty and C.Wang for animal care; P. Lovelace, S. Weber and C. Crumpton for FACS facility management. The authors would like to acknowledge ongoing support for this work: NIH/NCI Outstanding Investigator Award R35CA220434, NIH NIDDK R01DK115600, NIH NIAID R01AI143889 and Virginia and D.K. Ludwig Fund for Cancer Research (I.L.W.), American Society of Hematology (V.L.M.), NIA R36 award R36AG090859 (A.B), California Institute for Regenerative Medicine (K.T.), (A.W.) (R.Y.), (T.W.). H.N. was supported by the NIH (R01DK116944; R01HL147124), the Ludwig Foundation, and the Japan Society of the Promotion of Science. A.C.W. was supported by the NIH (K99HL150218) and the Leukemia and Lymphoma Society (3385-19).

## Author Contributions

V.L.M conceived, performed, analyzed and oversaw the experiments. K.T. and T.W. performed experiments under the supervision of V.L.M. A.W aided with design of initial flow cytometry experiments and data interpretation. V.L.M and I.L.W wrote the manuscript.

## Supplemental information titles and legends

**Supplemental Figure 1. Hoxb5 is expressed in lineage negative (Lin^-^) fetal liver compartments.** A. Flow cytometry plots of E14.5 fetal liver from mice depicting Hoxb5-tri-mCherry reporter activity in Hoxb5 high (Hoxb5^hi^), Hoxb5 low (Hoxb5^lo^), and Hoxb5 negative (Hoxb5^-^) populations in sequentially gated immunophenotype subpopulations: i. Lin^-^, ii. LKS, iii. LKS CD48^-^ FLK2^-^, iv. LKS CD48^-^ FLK2^-^CD150^+^ Mac1^+^; B. Frequency of each sequentially gated subpopulation in the parent compartment of the HSC immunophenotype profile at E12.5, E14.5 and E16.5; C. Frequency of Hoxb5^+^ cells (i) and count of Hoxb5^+^ cells (ii) in each sequentially gated subpopulation at E12.5, E14.5 and E16.5.

**Supplemental Figure 2. Donor peripheral blood chimerism by week and donor cell number.** A. Mean donor peripheral-blood chimerism at week 4-16 from 200, 100, 10, or 1 CD45.2^+^ Hoxb5^+^ FL-HSCs in primary and secondary recipients. Competitive transplantation against 1×10^6^ bone-marrow competitors. Individual points represent an individual mouse. B. Mean peripheral-blood chimerism of 10 Hoxb5+ or Hoxb5- FL-HSCs by competitive transplantation against 1×10^6^ CD45.1+ bone-marrow competitors represented as % CD45.2 chimerism. C. 10 cell donor multilineage chimerism in myeloid (GR1^+^ granulocytes and Mac1^+^ monocytes) and lymphoid (CD3^+^ T cells and B220^+^ B cells) populations.

**Supplemental Figure 3. Clustering and gene expression analysis of fetal liver HSCs. A.** UMAP of single-cell RNA sequencing data from fetal liver HSPCs showing Leiden clustering and annotation of cell types based on marker gene expression**. B.** UMAP showing unsupervised subclustering of fetal liver HSCs into two transcriptionally distinct clusters identified by Leiden algorithm.**C.** Feature plots showing normalized expression of representative genes enriched in cluster 1 (*Hoxb5⁺*-enriched). **D.** Feature plots showing normalized expression of representative genes enriched in cluster 0 (*Hoxb5⁻*-enriched).

**Supplemental Figure 4. Temporal gene expression in fetal liver HSCs by embryonic day.** Violin plots showing normalized expression of selected genes in *Hoxb5⁺* and *Hoxb5⁻* fetal liver HSCs at E12.5, E14.5, and E16.5.

**Supplemental Figure 5. Hoxb5 expression in E14.5 fetal liver hematopoietic stem and progenitor cell (HSPC) compartments.** A. Flow cytometry plots of E14.5 fetal liver from mice depicting Hoxb5-tri-mCherry reporter activity in HSPC immunophenotype subpopulations including: HSC (Lin^-^ Sca^+^ cKit^+^ CD150^+^ FLK2^-^ CD34^-^); MPPb (Lin^-^ Sca^+^ cKit^+^ CD150^-^ FLK2^-^ CD34^+^); MPPc (Lin^-^ Sca^+^ cKit^+^ CD150^-^ FLK2^+^ CD34^+^); CMP (Lin^-^ Sca^lo/-^ cKit^+^ CD34^med/hi^ CD16/32^-/lo^); MEP (Lin^-^ Sca^lo/-^ cKit^+^ CD34^-^CD16/32^-/lo^ CD150^+^); GMP (Lin^-^ Sca^lo/-^ cKit^+^ CD34^hi^ CD16/32^hi^; B. Graphical representation of Hoxb5 expression in HSPC compartments expressed as: C. Hoxb5^+^ cells in each HSPC compartment as a frequency of Lin^-^ fetal liver cells, E. Hoxb5^+^ cell frequency in each HSPC compartment, and D. Hoxb5^+^ cell count for each HSPC compartment per 10000 Lin- cells.

**Supplemental Figure 6.** A. UMAP of single-cell transcriptomes from fetal liver HSPCs showing Leiden clustering and annotation of major progenitor populations (HSC, HSPC, CMP, GMP, MEP). B. UMAP of all fetal liver HSPCs colored by developmental stage (E12.5, E14.5, E16.5). C. UMAP displaying *Hoxb5-mCherry* reporter status (red, *Hoxb5-mCherry*⁺; gray, *Hoxb5-mCherry*⁻) and bar graph showing the proportion of reporter-positive cells within each population. D. Dot plot showing expression of canonical HSC and progenitor marker genes across annotated populations. E. Feature and violin plots showing normalized expression of *Hoxb5*, *Hlf*, and *Mecom* across HSC, HSPC, CMP, GMP, and MEP clusters.

## Methods

### Mice

Mice were bred at Stanford University animal facility according to NIH guidelines. All animal protocols were approved by the Stanford University Administrative Panel on Laboratory Animal Care. C57BL/6J tri-mCherry mice (CD45.2^+^) were bred for timed pregnancy. C57BL/6J timed pregnant mice (Jackson Laboratory) were used as wild-type controls. Ten-to-twelve-week-old B6.SJL-*Ptprc^a^ Pepc^b^*/BoyJ female mice (Jackson Laboratory) were used as recipients for transplantation assays. Supporting cells for competitive reconstitution assays were collected from B6.SJL-*Ptprc^a^ Pepc^b^*/BoyJ (CD45.1^+^).

### Fluorescent activated cell sorting of stem and progenitor cells

Fetal liver cells were obtained from timed pregnant C57BL6/J tri-mCherry Hoxb5 reporter (CD45.2^+^) and control C57BL6/J mice at embryonic day (E) 12.5, E14.5 and E16.5. Fetal liver cells were dissociated by manual homogenization in Ca^2+^ and Mg^2+^ free PBS supplemented with 2% heat inactivated bovine serum (Gibco). For mouse bone-marrow cells were isolated from the tibia, femur and pelvis. Cells were passed through 40µm strainers before analysis and sorting. To enrich for HSCs, cells were stained with APC-conjugated anti-c-Kit (2B8) and fractionated using Anti-APC magnetic beads and LS columns (Miltenyi Biotec). Cells were stained with conjugated antibodies to the surface markers: c-Kit, Sca-1, Flk2, CD48, CD150, and Mac1 and conjugated lineage markers including Ter119, CD3, B220, CD4, CD8, and GR1. For lymphoid populations, fetal liver cells were stained with antibodies against B220 and CD3. For myeloid populations, cells were stained with antibodies against Gr1 and Mac1. Antibody staining was performed at 4C and cells were incubated for 30 minutes. Cells stained with CD34 were incubated for 90 minutes at 4C. Before sorting cells were stained with DAPI to assess viability as per the manufacturers recommendations. Flow cytometry and cell sorting were performed on a FACS Aria II cell sorter (BD Biosciences) and analyzed using FlowJo software (Tree Star).

### Competitive Transplant Assays

Adult B6.SJL-*Ptprc^a^ Pepc^b^/BoyJ* (Jackson Laboratory) recipients (10-12 weeks old) were lethally irradiated at a single dose of 9.5 Gy. For reconstitution assays, CD45.2^+^ candidate donor cells were sorted and transplanted with 2x10^5^ CD45.1^+^ whole bone marrow cells into the retro-orbital venous sinus or the tail vein of anesthetized CD45.1^+^ recipients, following lethal irradiation. Donor chimerism was tracked by collecting peripheral blood from recipient CD45.1 mice at 4, 8, 12, and 16 weeks after primary and secondary transplantations. Peripheral blood was subject to ammonium-chloride-potassium (ACK) cell lysis and subsequently analyzed to determine the level and hematopoietic lineage of the donor reconstitution by staining with anti-CD45.1, anti-CD45.2, anti-CD11b, anti-LY-6G/LY-6C, anti-CD45R, anti-CD3 and anti-B220 antibodies. Following a wash step, cells were analysed by flow cytometry using propidium iodide as a dead stain. Secondary bone-marrow transplantation assays were performed by transferring 1×10^6^ bone-marrow cells from the primary recipient mice into lethally irradiated CD45.1 secondary recipient mice, with donor chimerism analysed as above.

### Frequency of donor cell engraftment

Set numbers of CD45.2^+^ donor cells were aliquoted by fluorescence-activated cell sorting (FACS) as above, and transplanted into lethally irradiated CD45.1^+^ recipients, transplanted with 2x10^5^ CD45.1^+^ whole bone marrow competitor cells. Donor chimerism was analysed as above. Mice showing long-term (> 16 week) > 1% peripheral blood multi-lineage chimerism (at least 0.2% myeloid donor chimerism) counted as the threshold for positive engraftment. Engraftment frequencies were estimated using a single-hit Poisson model. By performing single-cell transplants, the engraftment probability can be estimated directly as a binomial proportion with confidence intervals. Frequencies were expressed to indicate the number of phenotypically defined cells containing one functional long-term HSC. Frequency of donor cell engraftment was calculated using the Extreme Limiting Dilution Analysis (ELDA) software (Hu and Smyth, 2009; https://bioinf.wehi.edu.au/software/elda/).

### Single-cell RNA sequencing

Hematopoietic stem and progenitor cells (HSPCs) from mouse fetal liver were prepared as single-cell suspensions and sorted by flow cytometry as described above. Individual cells were index-sorted into 96-well plates containing lysis buffer following the general approach of Liu et al. Sorted plates were centrifuged at 4 °C, snap-frozen on dry ice, and stored at −80 °C until processing. Reverse transcription and cDNA pre-amplification were performed according to the Smart-seq3 protocol with minor modifications. For each well, 1 μL of purified cDNA was assessed by Fragment Analyzer (Advanced Analytical) for concentration and fragment size distribution. Wells yielding less than 1.7 ng/μL cDNA were excluded; this threshold was determined from negative control wells containing ERCC spike-ins but no sorted cell. cDNA passing quality control was reformatted into 384-well plates using a Mosquito X1 liquid handler (SPT Labtech) and normalized to 1.7–4.0 ng/μL using UltraPure water. Normalized cDNA was tagmented by combining 0.4 μL cDNA with 1.2 μL homebrew Tn5 transposase mix (1 ng/μL Tn5, 16 mM Tris-HCl pH 7.6, 16 mM MgCl₂, and 8% dimethylformamide). Reactions were stopped with 0.4 μL 0.1% SDS. Libraries were indexed by PCR using 5 μM i5 and i7 primers (Integrated DNA Technologies; 7680-plex unique dual indices) and KAPA HiFi HotStart ReadyMix, with the following thermocycling program: 72 °C 3 min; 95 °C 30 s; then 10 cycles of 98 °C 10 s, 67 °C 30 s, and 72 °C 60 s. One microliter from each well was pooled per 384-well plate and purified with 0.8× AMPure XP beads. Plate-level pools were quantified by Fragment Analyzer, normalized, and combined into 7680-cell libraries, which were again purified and concentrated using 0.8× AMPure beads. Final pooled libraries were sequenced on an Illumina NovaSeq 6000 S4 flow cell to generate ∼1–2 million paired-end 150 bp reads per cell. Demultiplexing was performed using bcl2fastq v2.19. Adapter trimming was done with Skewer v0.2.2. Reads were aligned to the mouse reference genome (GRCm38.p6, Gencode vM25) using STAR v2.6.1d with a two-pass strategy. In the first pass, splice junctions detected across cells were aggregated and incorporated into a new STAR genome index, which was then used for second-pass mapping. Alignment parameters followed ENCODE long-mRNA pipeline recommendations, with “--quantMode TranscriptomeSAM” enabled to produce transcriptome BAM files. Gene- and transcript-level quantifications were generated using RSEM v1.3.3 with the “--single-cell-prior” option to account for sparse transcript detection typical of single-cell data. Gene count matrices were processed in Scanpy v1.8.2. Genes detected in fewer than three cells and cells expressing fewer than 500 genes or 5,000 total UMI counts were removed. Counts were normalized to 10,000 reads per cell, log₁₀-transformed, and scaled to a maximum value of 10. Highly variable genes were identified using Scanpy default parameters. Principal component analysis (PCA) was performed, followed by nearest-neighbor graph construction and Leiden clustering. Data visualization and lineage trajectory reconstruction were conducted using UMAP and PAGA implementations within Scanpy.

### Genotyping

Genomic DNA from Hoxb5-trimCherry mice was isolated from tail biopsies using QuickExtract solution (EpiCentre). PCR amplification was performed using forward primer ( 5′-GACGTATCGAGATCGC CCAC-3′) and reverse primer (5′-CCTTGGTCACCTTCAGCTTGG-3′).

### Statistics

Comparative analyses were performed using unpaired Student’s t tests or One way ANOVA. Pearson’s chi-squared test was performed using online software (http://vassarstats.net).

