## Supplemental Figures for "Hoxb5 Enriches Long-Term Hematopoietic Stem Cell Activity within the Mouse Fetal Liver Phenotypic HSC Compartment"

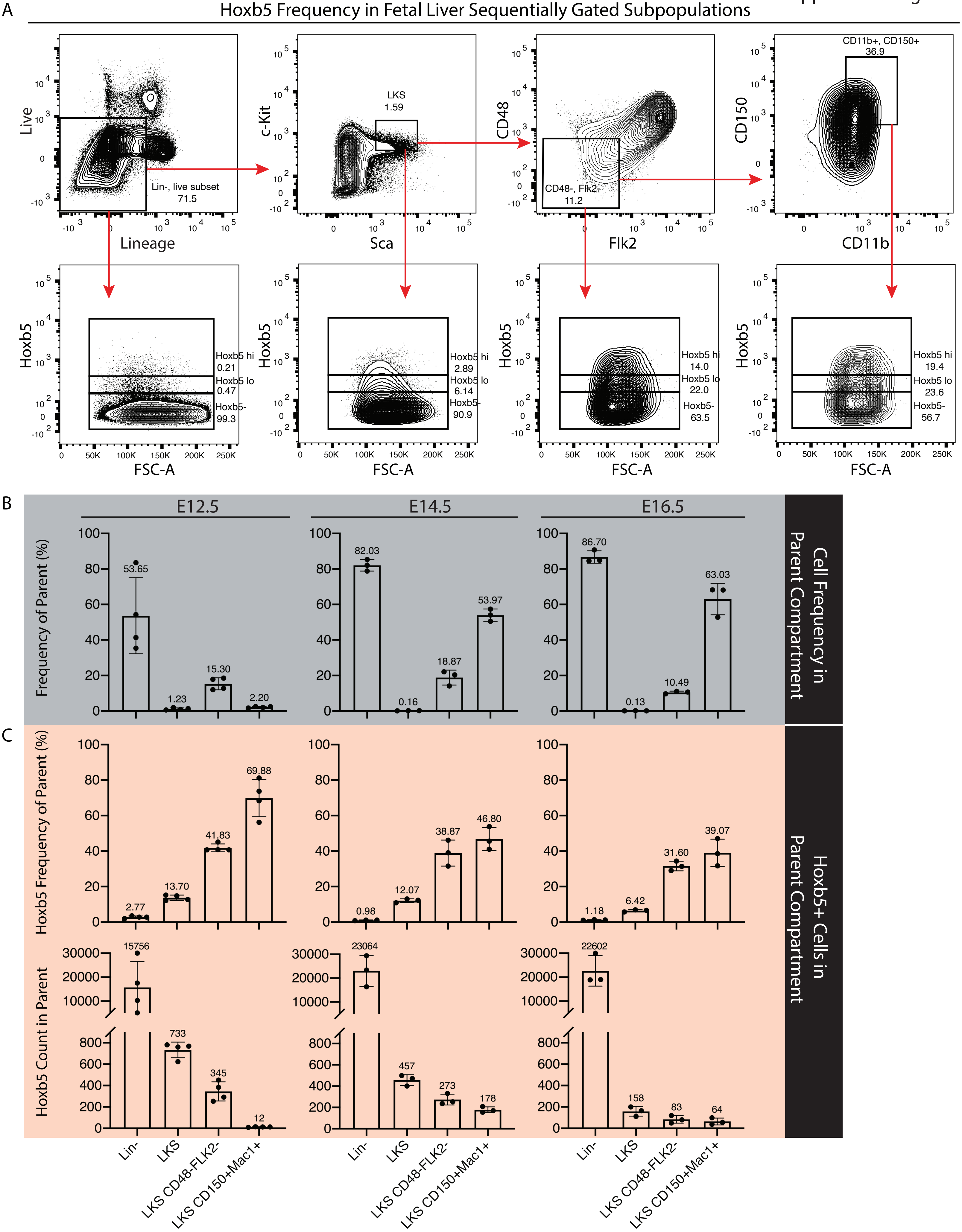
A
Hoxb5 Frequency in Fetal Liver Sequentially Gated Subpopulations
Live
Lineage
Lin-, live subset 71.5
c-Kit
Sca
LKS 1.59
CD48
Flk2
CD48-, Flk2- 11.2
CD150
CD11b
CD11b+, CD150+ 36.9
Hoxb5
FSC-A
Hoxb5 hi 0.21
Hoxb5 lo 0.47
Hoxb5- 99.3
Hoxb5 hi 2.89
Hoxb5 lo 6.14
Hoxb5- 90.9
Hoxb5 hi 14.0
Hoxb5 lo 22.0
Hoxb5- 63.5
Hoxb5 hi 19.4
Hoxb5 lo 23.6
Hoxb5- 56.7
B
E12.5
E14.5
E16.5
Frequency of Parent (%)
53.65
1.23
15.30
2.20
82.03
0.16
18.87
53.97
86.70
0.13
10.49
63.03
Cell Frequency in Parent Compartment
C
Hoxb5 Frequency of Parent (%)
2.77
13.70
41.83
69.88
0.98
12.07
38.87
46.80
1.18
6.42
31.60
39.07
Hoxb5 Count in Parent
15756
733
345
12
23064
457
273
178
22602
158
83
64
Hoxb5+ Cells in Parent Compartment
Lin-
LKS
LKS CD48-FLK2-
LKS CD150+Mac1+

A Mean Donor Peripheral Blood Chimerism Week 4-16 from Hoxb5$^{+}$ HSCs

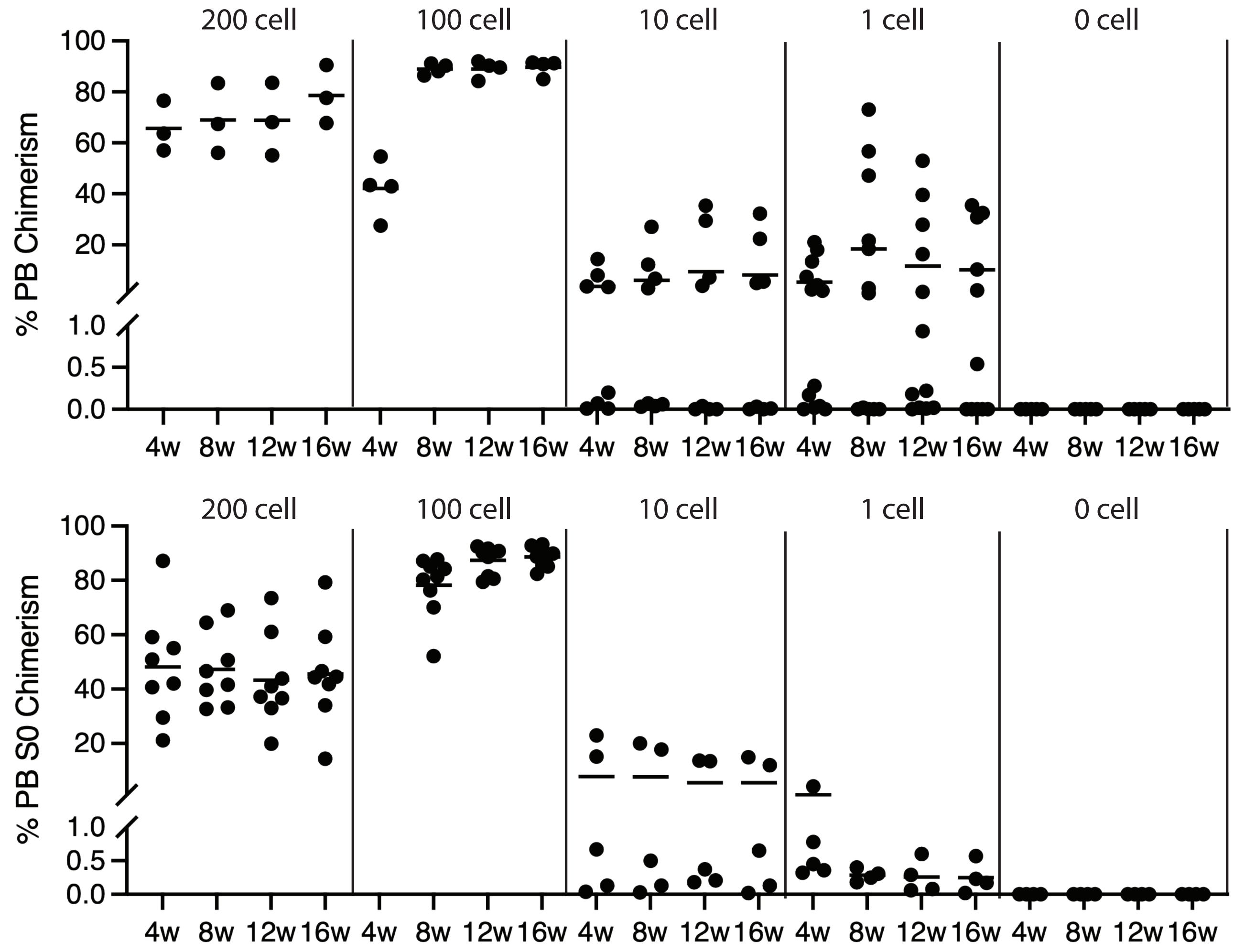


10 Cell Peripheral Blood Chimerism Week 4 -16

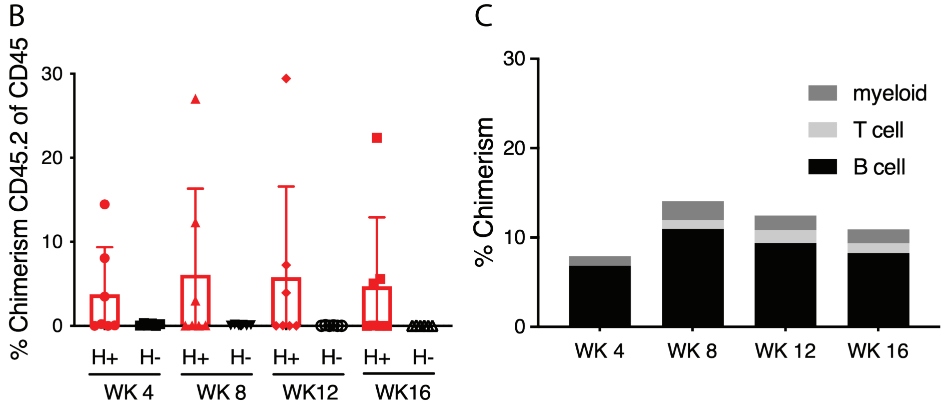

Supplemental Figure 3

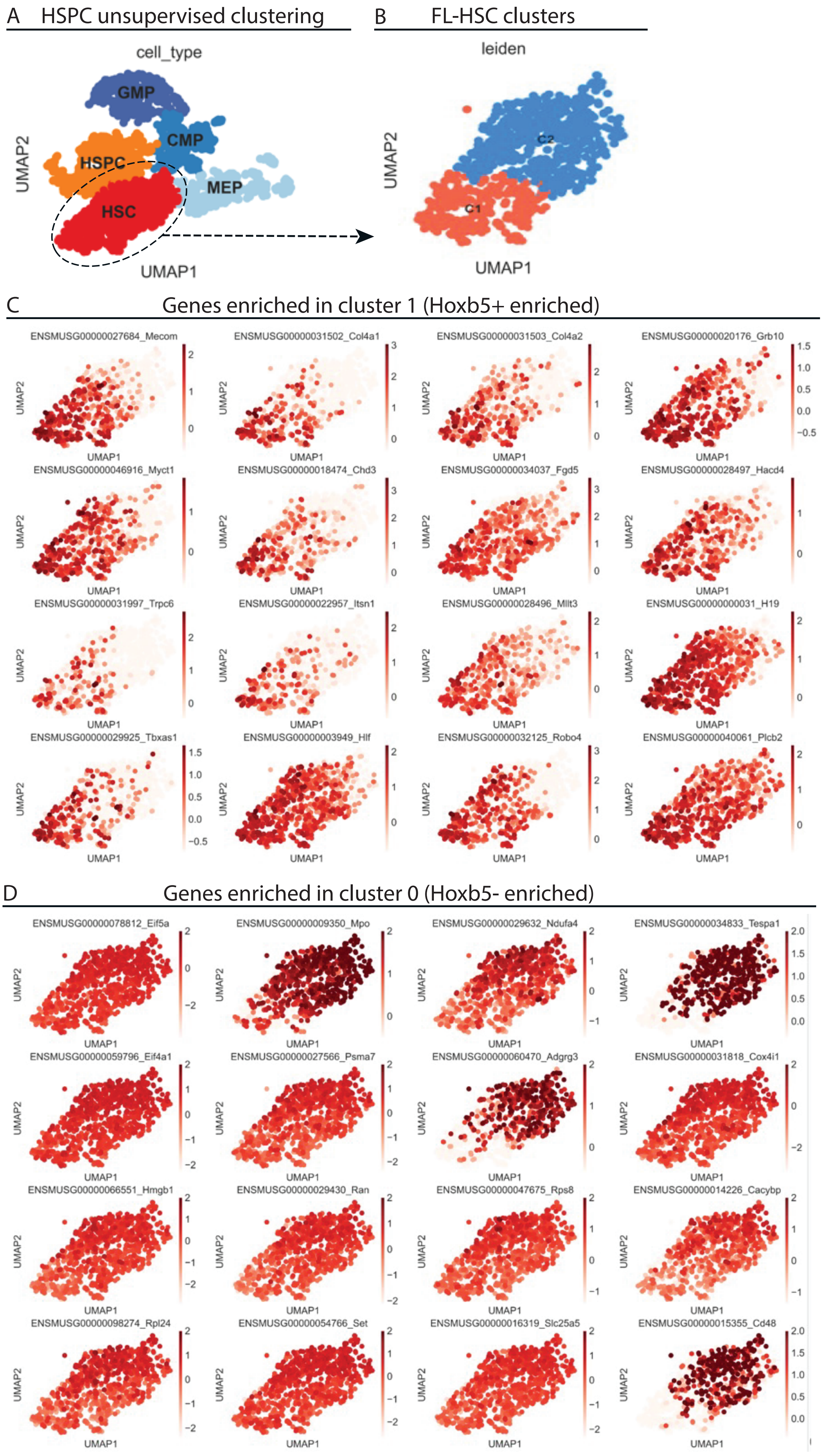

Supplemental Figure 4

Temporal gene expression in FL-HSCs by embryonic day

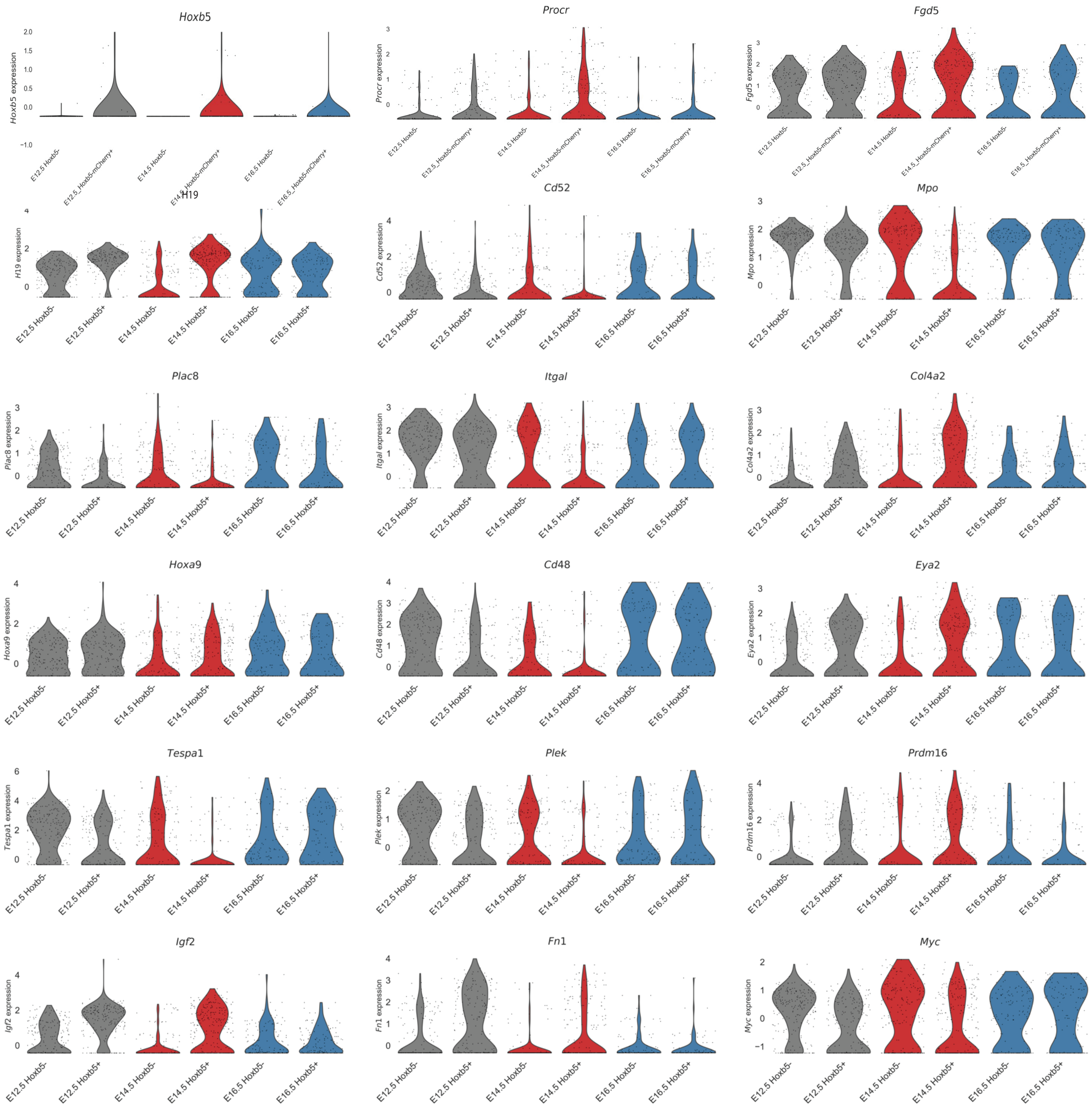

Supplemental Figure 5

A

### Hoxb5 is expressed in Fetal Liver HSPCs

| | |
|---|---|
| HSC | Lin⁻ cKit⁺ Sca1⁺ CD150⁺ Flk2⁻ CD34⁺ |
| MPPa | Lin⁻ cKit⁺ Sca1⁺ CD150⁺ Flk2⁻ CD34⁺ |
| MPPb | Lin⁻ cKit⁺ Sca1⁺ CD150⁻ Flk2⁻ CD34⁺ |
| MPPc | Lin⁻ cKit⁺ Sca1⁺ CD150⁻ Flk2⁺ CD34⁺ |
| CMP | Lin⁻ cKit⁺ Sca1$^{lo/-}$ CD34$^{med/hi}$ CD16/32$^{-/lo}$ |
| MEP | Lin⁻ cKit⁺ Sca1$^{lo/-}$ CD34⁻ CD16/32$^{-/lo}$ CD150⁺ |
| GMP | Lin⁻ cKit⁺ Sca1$^{lo/-}$ CD34$^{hi}$ CD16/32$^{hi}$ |

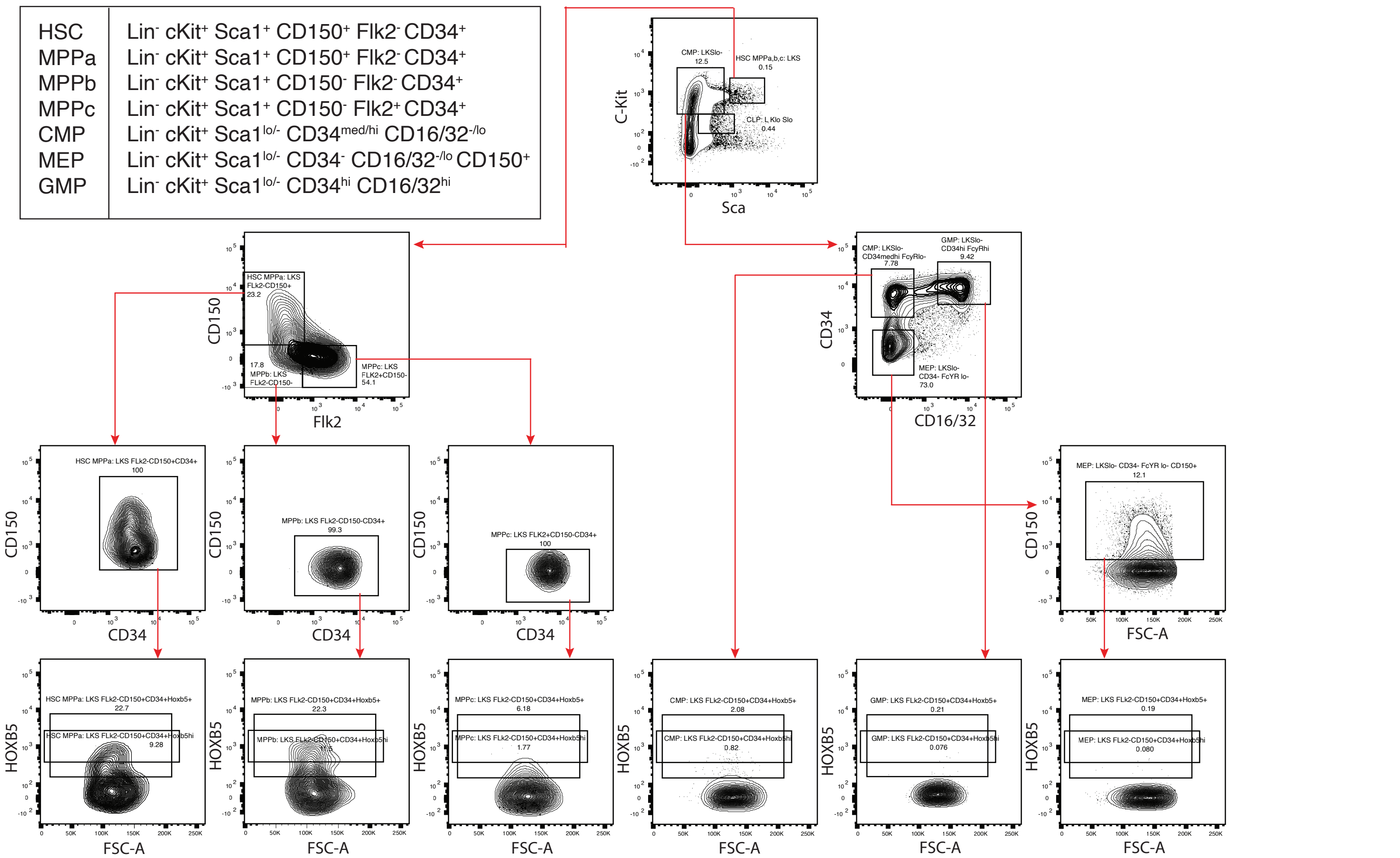


### Hoxb5 Expression in Fetal Liver HSPCS

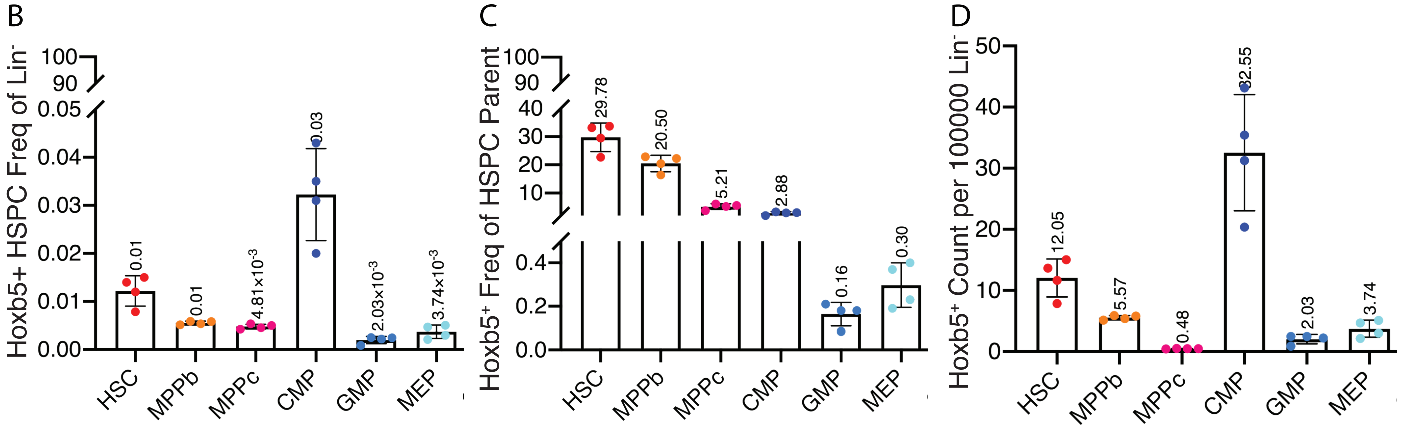

### Supplemental Figure 6

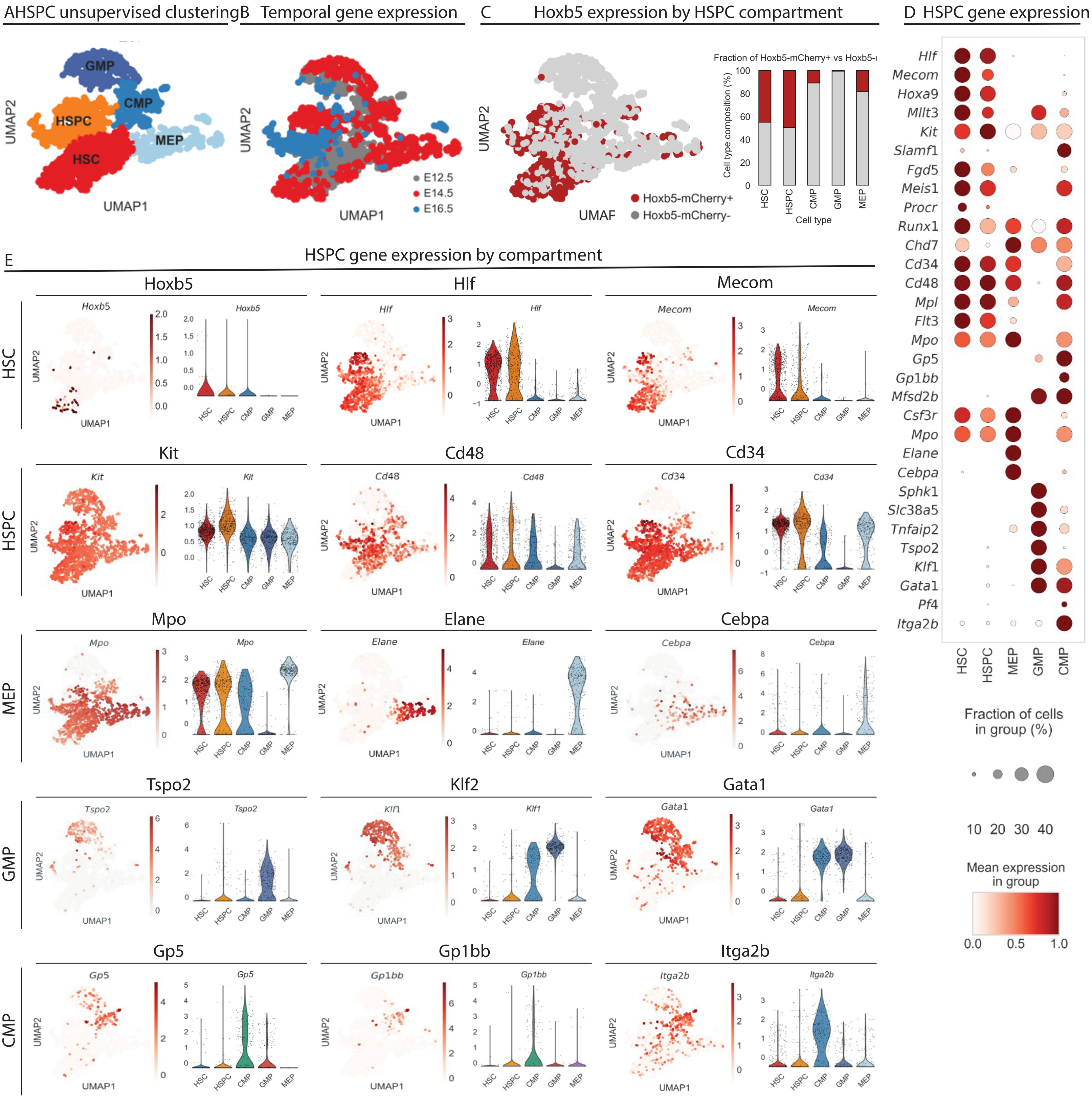
